# Transcriptional suppression of sphingolipid catabolism controls pathogen resistance in *C. elegans*

**DOI:** 10.1101/2023.08.10.552843

**Authors:** Mohamad A. Nasrallah, Nicholas D. Peterson, J. Elizabeth Salisbury, Pengpeng Liu, Amanda L. Page, Samantha Y. Tse, Khursheed A. Wani, Claire E. Tocheny, Read Pukkila-Worley

## Abstract

Sphingolipids are required for diverse biological functions and are degraded by specific catabolic enzymes. However, the mechanisms that regulate sphingolipid catabolism are not known. Here we characterize a transcriptional axis that regulates sphingolipid breakdown to control resistance against bacterial infection. From an RNAi screen for transcriptional regulators of pathogen resistance in the nematode *C. elegans*, we identified the nuclear hormone receptor *nhr-66,* a ligand-gated transcription factor homologous to human hepatocyte nuclear factor 4. Tandem chromatin immunoprecipitation-sequencing and RNA sequencing experiments revealed that NHR-66 is a transcriptional repressor, which directly targets sphingolipid catabolism genes. Transcriptional de-repression of two sphingolipid catabolic enzymes in *nhr-66* loss-of-function mutants drives the breakdown of sphingolipids, which enhances host susceptibility to infection with the bacterial pathogen *Pseudomonas aeruginosa*. These data define transcriptional control of sphingolipid catabolism in the regulation of cellular sphingolipids, a process that is necessary for pathogen resistance.

## INTRODUCTION

Sphingolipids are complex amphipathic macromolecules that are required for the integrity, fluidity, and barrier function of cell membranes. Sphingolipids are also metabolized to bioactive signaling molecules, such as sphingosine-1-phosphate, that have been implicated in fibrosis, cell proliferation, neurotransmission, and immune-cell trafficking^1–8^. In addition, ceramides, a sub-class of sphingolipids, are important for apoptosis, response to cellular stress, and innate immunity^9–16^. Sphingolipid levels are therefore tightly regulated by specific biosynthetic and catabolic pathways.

Ceramides and other sphingolipids are broken down in cells by specific enzymes. For example, acid ceramidase catabolizes ceramides to sphingosines, which can be recycled back to ceramides or phosphorylated to produce bioactive lipids^17^. The final exit point of sphingolipid catabolism is the irreversible breakdown of sphingosine-1-phosphate into long-chain fatty acid aldehyde and phospho-ethanolamine by sphingosine phosphate lyase^2^. Importantly, several human inborn errors of metabolism, such as Tay-Sachs and Niemann-Pick diseases, are caused by defects in the breakdown of sphingolipids^18–20^. Although the biochemical steps in sphingolipid and ceramide catabolism are defined, it is not known how these processes are regulated and whether such mechanisms contribute to stress resistance.

Gene regulatory networks sense metabolic perturbations in the cell and regulate the flux of cellular metabolites by changing the transcription of specific enzymes^21,22^. A few specific examples of such metabolic gene regulatory networks have been characterized, particularly in the regulation of specific metabolites, lipids, and cholesterol, which promote adaptation to diet or changing environmental conditions^23–30^. Many of these networks utilize nuclear hormone receptors (NHRs), which are ligand-gated transcription factors that regulate cellular physiology by controlling gene expression. The nematode *Caenorhabditis elegans* has been particularly useful for characterizing metabolic gene regulatory networks, in part because its genome encodes a large number of *nhr* genes compared to other metazoans. Nematodes express 274 NHRs compared to 42 in humans and 24 in *Drosophila,* the great majority of which have unknown functions^31–35^. Intriguingly, of the few *C. elegans* NHRs that have been characterized in detail, several have specialized functions in pathogen detection or immunometabolism. For example, one *C. elegans* NHR, NHR-86, surveys for a pathogen-derived metabolite — a “pattern of pathogenesis” specifically associated with infection or bacterial virulence — to assess the relative threat of a bacterial pathogen in its environment and activate host immune defenses^36^. Other *C. elegans nhr* genes regulate metabolism to promote survival during pathogen infection. The *C. elegans* peroxisome proliferator-activated receptor ⍺ (PPAR⍺) homolog NHR-49 is a master regulator of lipid metabolism and *nhr-49* mutants are dramatically hypersusceptible to killing by bacterial pathogens^37,38^. Likewise, the monounsaturated fatty acid oleate and the methyl donor S-adenosylmethionine, whose levels are each regulated by NHR-49, are individually necessary to survive pathogen challenge^39,40^. In addition, control of cellular cholesterol levels by NHR-8, a *C. elegans* homolog of mammalian liver X receptor and pregnane X receptor, primes protective activation of the p38 PMK-1 innate immune pathway^41^ and is required for pathogen resistance^41,42^. Nuclear hormone receptors also suppress the transcription of innate immune defense genes to promote immune homeostasis, both downstream of a key defense regulator^43^ and via a response linked to iron uptake^44^. Together, these studies characterize novel mechanisms of immunometabolism^45^ and suggest that the NHR family in *C. elegans* may have expanded, at least in part, because of their diverse roles in promoting survival during pathogen infection.

Here, we demonstrated that regulation of sphingolipid breakdown occurs at the transcriptional level and showed that this process is required for pathogen resistance in *C. elegans*. We discovered that NHR-66 suppresses sphingolipid catabolism by directly repressing the transcription of two enzymes: sphingosine phosphate lyase (*spl-2*) and acid ceramidase (*asah-2*). Transcriptional de-repression of *spl-2* and *asah-2* in *nhr-66* loss-of-function mutants drove the excessive breakdown of sphingolipids and ceramides, which compromised host survival during pathogen infection. These data demonstrated for the first time that transcriptional regulation of sphingolipid catabolism is required for pathogen resistance, revealing a novel pathway that can be targeted to promote survival during bacterial infection.

## RESULTS

### An RNAi screen identifies nuclear hormone receptors required for pathogen resistance in *C. elegans*

We performed an RNAi screen to identify the *C. elegans nhr* genes required for host survival during bacterial infection (Fig. 1A). From a primary screen of 271 of the 274 *nhr* genes in the *C. elegans* genome, we identified 15 *nhr* genes that, when knocked down, reduced *C. elegans* survival during infection with the bacterial pathogen *P. aeruginosa* (Figs. 1A–C, Table S1). Lifespan assays, performed by monitoring survival of nematodes fed on a standard bacterial diet (*E. coli* OP50), revealed that 10 of these 15 *nhr* genes affected pathogen susceptibility, but without pleiotropic effects on nematode lifespan (Figs. 1B and 1C, Table S1). We then performed three independent trials of *P. aeruginosa* pathogenesis assays in *C. elegans* following RNAi-mediated knockdown of these 10 *nhr* genes (Fig. S1A, Table S1). RNAi targeting four of these *nhr* genes (*nhr-8*, *nhr-66*, *nhr-31,* and *nhr-32*) rendered *C. elegans* hypersusceptible to pathogen-mediated killing in three of three biological replicate trials (Fig. S1A, Table S1).

**Figure 1.**
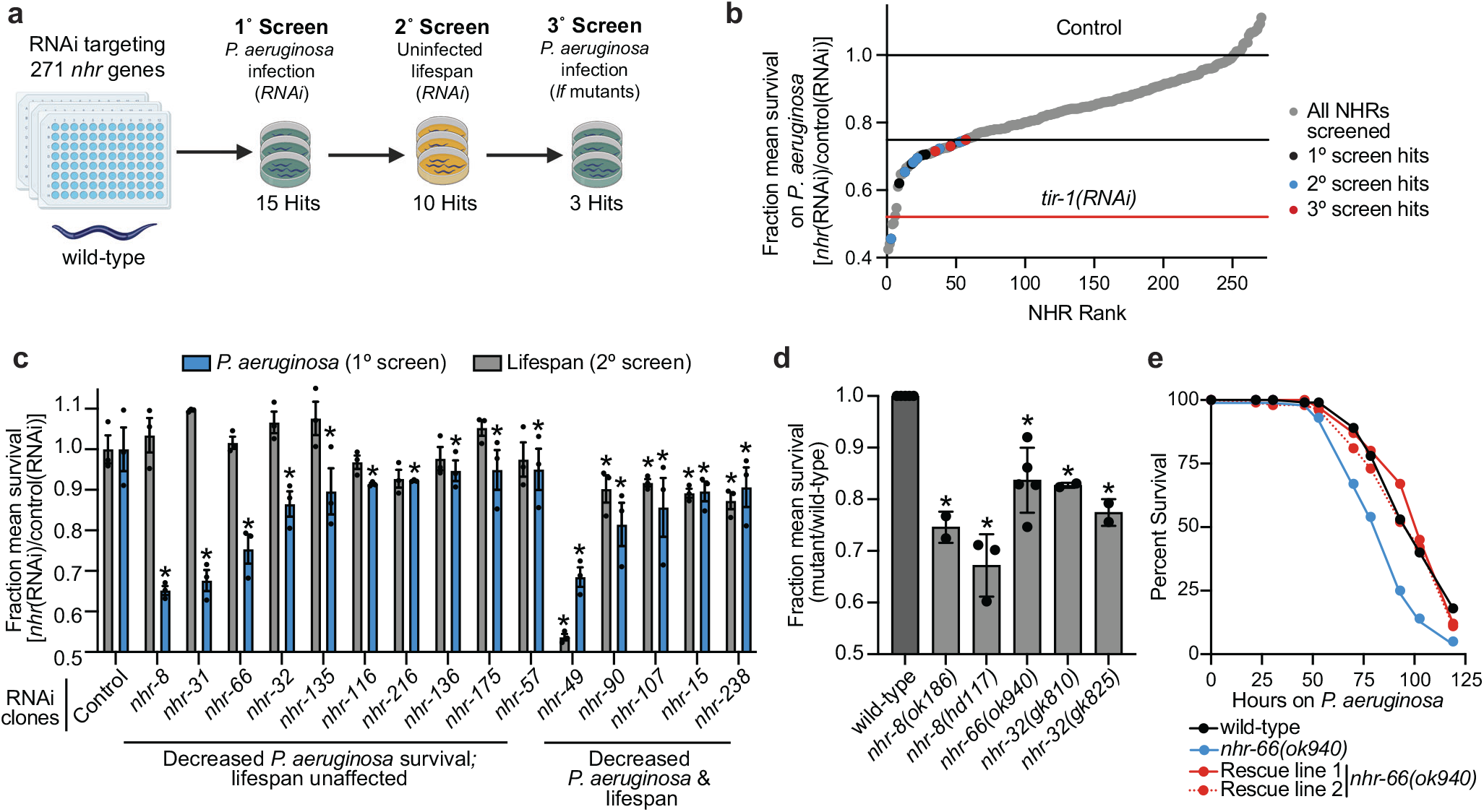
An RNAi screen identifies nuclear hormone receptors required for pathogen resistance in *C. elegans*. **A.** A schematic of the RNAi screen for 271 of the 274 *nhr* genes that control resistance to *P. aeruginosa* infection in *C. elegans* is shown. **B.** The fraction mean survival compared to wild-type animals on *P. aeruginosa* is plotted for each of the *C. elegans nhr* RNAi clones in the screen. Hits from the primary, secondary, and tertiary screens are indicated. The fraction mean survival for the control *tir-1(RNAi)* is shown. **C.** The fraction mean survival during *P. aeruginosa* infection and in a lifespan assay is compared for each of the 15 *nhr* genes that were identified in the primary screen. Data are the average of three replicate trials with error bars showing SEM (n=3) with error bars representing SEM. * equals p<0.05 (Kaplan-Meier method with log-rank test). **D.** The fraction mean survival during *P. aeruginosa* infection was plotted for the indicated mutant genotypes. Mean survival is the average of 2-5 biological replicates and is expressed relative to the survival of wild-type *C. elegans*, with error bars representing SEM. * equals p<0.05 (Kaplan-Meier method with log-rank test). **E**. *C. elegans*-*P. aeruginosa* pathogenesis assay with *C. elegans* of the indicated genotypes. The difference in survival between *nhr-66(ok940)* and the other genotypes is significant (*p*<0.05, log-rank test, n=3). Data shown are representative of three independent trials. Sample sizes, mean lifespan, and p-values for each trial are shown in Table S5. The source data for the RNAi screen is in Table S1. See also Figure S1.

Loss-of-function mutants in three of these four *nhr* genes died significantly faster than wild-type animals during bacterial infection (Figs. 1D and 1E, Figs. S1B and S1C). The *nhr-31(tm1547)* mutant is embryonically lethal^46^. We confirmed that mutations at these specific loci led to pathogen hypersusceptibility by re-introducing a wild-type copy of each of these *nhr* genes, expressed under the control of their own promoters, into the *nhr-8(hd117)*, *nhr-32(gk810)*, and *nhr-66(ok940)*, loss-of-function mutant backgrounds. Complementation of *nhr-8* (Fig. S1D)^42^, *nhr-32* (Fig. S1E), and *nhr-66* (Fig. 1E) in this manner restored wild-type resistance to pathogen infection. Thus, *nhr-8*, *nhr-32*, and *nhr-66* specifically and robustly enhance *C. elegans* survival during *P. aeruginosa* infection.

The recovery of *nhr-8* provided an internal control for the RNAi screen, as we previously characterized its role in immune regulation and pathogen resistance^41^. Likewise, *nhr-49* is required for normal lifespan in *C. elegans* and resistance to pathogen infection^37,38,47,48^, findings we also observed (Fig. 1C). We chose 12 *nhr* genes that were not recovered in our primary screen and confirmed that knockdown of these genes did not affect resistance to *P. aeruginosa* infection (Fig. S1F). Together, these results validated the RNAi screen, which identified specific *C. elegans nhr* genes that are required for resistance to *P. aeruginosa* infection.

### NHR-66 is a transcriptional repressor that directly targets stress response and sphingolipid metabolism genes

To further characterize the role of NHR-66 in promoting resistance to pathogen infection, we performed chromatin immunoprecipitation sequencing (ChIP-seq) and RNA sequencing (RNA-seq). For these experiments, we used CRISPR-Cas9 to insert a 3xFLAG sequence at the *nhr-66* locus (Fig. S2A). NHR-66 protein-DNA complexes were precipitated using an anti-FLAG antibody and deep sequenced. In parallel, mRNA transcripts were quantified in *nhr-66*(*ok940*) mutant and wild-type animals. A comparison of the ChIP-seq and RNA-seq datasets revealed that NHR-66 bound to the promoters and regulated the transcription of 18 genes (Fig. 2A–E, Table S2, and Table S3). We validated the ChIP-seq data using PCR to determine enrichment of promoter sequences of three NHR-66 targets [*spl-2* (Fig. 2F), *asah-2* (Fig. 2G), and *cyp-13A4* (Fig. 2H)] from biological replicate ChIP samples (ChIP-PCR). In addition, immunoprecipitation with an anti-FLAG antibody did not enrich for random intergenic regions in NHR-66:FLAG animals, as measured by ChIP-seq (Fig. 2I) or ChIP-PCR (Figs. 2J and 2K). The RNA-seq dataset was validated using qRT-PCR to measure the expression of *spl-2* (Fig. S2B), *asah-2* (Fig. S2C), and *cyp-13A4* (Fig. S2D).

**Figure 2.**
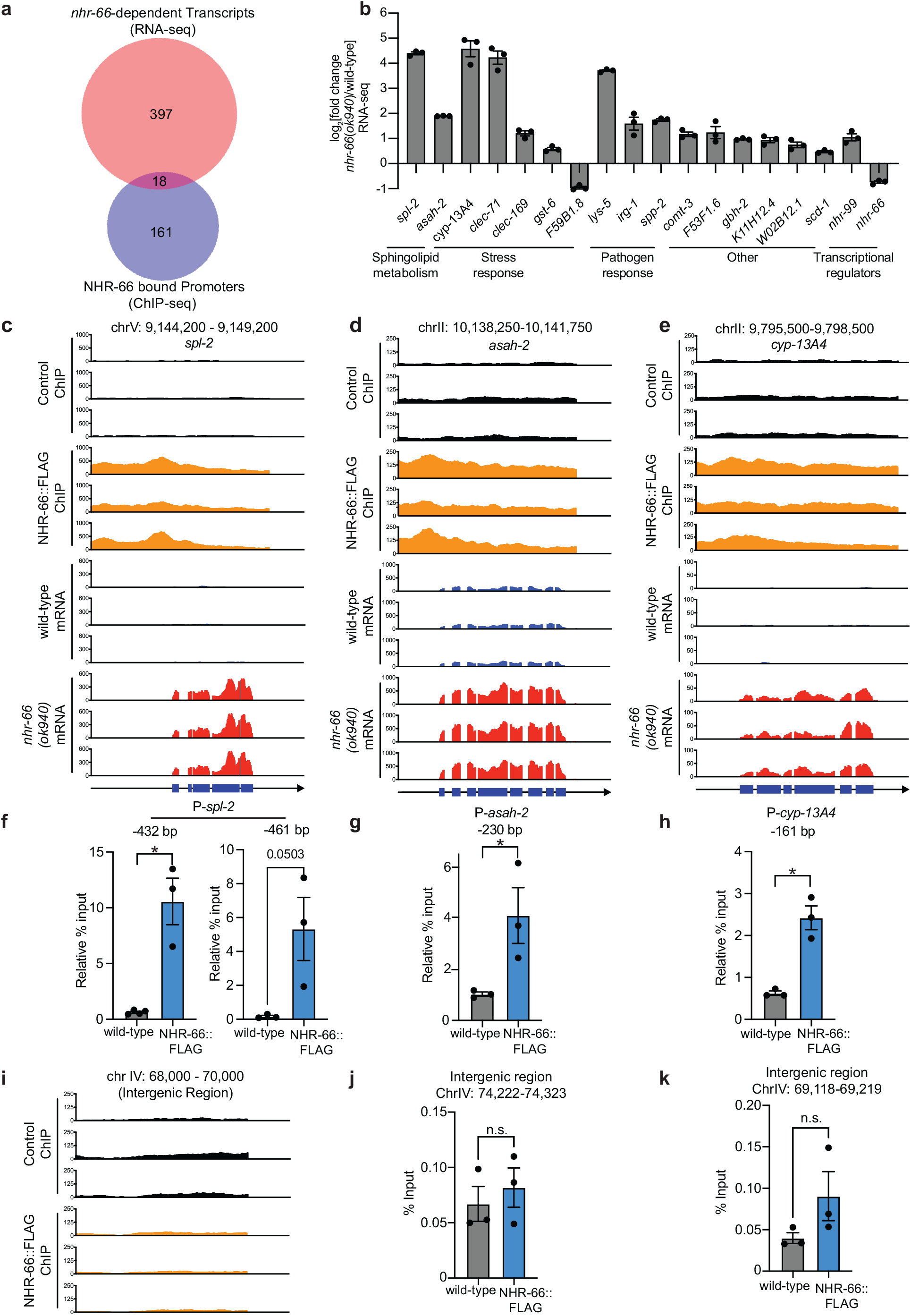
NHR-66 is a transcriptional repressor that directly targets stress response and sphingolipid metabolism genes. **A.** Venn diagram showing the overlap between the genes whose promoters were bound by NHR-66::FLAG in the ChIP-seq experiment and the genes that are differentially regulated in *nhr-66(ok940)* mutants compared to wild-type (from the RNA-seq experiment). The overlap between these datasets is significant (hypergeometric *p*-value = 1.42 x 10^-3^). For both the RNA-seq and ChIP-seq experiments, n=3 biological replicates, q<0.01. **B.** Fold change in expression of the 18 direct target genes of NHR-66 in *nhr-66*(*ok940*) mutants compared to wild-type animals is shown from the RNA-seq data. n=3 biological replicates, q<0.01. **C–E.** ChIP-seq and mRNA-seq profiles from each of the three biological replicates are presented for *spl-2* (C), *asah-2* (D), and *cyp-13A4* (E). The y-axis is the number of reads (log_2_). A gene model shows the location of the exons (blue) of the indicated genes. **F and G.** ChIP-PCR, performed to confirm the ChIP-seq data, is shown. The % input relative to the abundance of a random intergenic region of chromosome IV is presented for the promoter regions of *spl-2* (F) (two different regions are shown), *asah-2* (G) and *cyp-13A4* (H) (locations are indicated relative to the start codon of the gene). ChIP-seq profiles **(I)** and ChIP-PCR data **(J and K)** are shown for random intergenic regions on chromosome IV, as described above. For the ChIP-PCR data in this figure, n=3 biological replicates, * *p*<0.01 (Student’s unpaired t-test). Source data for this figure is in Table S6. See also Table S2 for the RNA-seq data, Table S3 for the ChIP-seq data, and Figure S2.

Interestingly, 16 of the 18 NHR-66 targets were expressed at higher levels in *nhr-66(ok940)* mutants than in wild-type (Fig. 2B). Thus, NHR-66 is predominantly a transcriptional repressor. In addition, NHR-66 directly promotes its own transcription, suggesting that it functions in a feedback loop (Fig. 2B). Two of the 18 NHR-66 transcriptional targets, *spl-2* and *asah-2,* function in sphingolipid catabolism^49^ and eight are stress or pathogen response genes. Thus, NHR-66 binds to the promoters of stress response and sphingolipid metabolism genes to repress their transcription.

### NHR-66 suppresses intestinal sphingolipid catabolism genes to promote resistance to *P. aeruginosa* infection

Given that both *spl-2* and *asah-2* are transcriptionally suppressed by NHR-66, we hypothesized that regulation of sphingolipid breakdown by NHR-66 is required for pathogen resistance in *C. elegans* and that hyperactivation of these enzymes accounts for the hypersusceptibility of *nhr-66(ok940)* mutants to pathogen infection. To test this hypothesis, we used RNAi to knockdown the NHR-66 target genes *spl-2* and *asah-2* in the *nhr-66(ok940)* mutant background. Importantly, RNAi of both *spl-2* (Figs. 3A and 3B) and *asah-2* (Figs. 3C and 3D) restored wild-type susceptibility to pathogen infection. qRT-PCR of *spl-2* (Fig. 3B) and *asah-2* (Fig. 3D) confirmed that RNAi depleted these transcripts in the *nhr-66(ok940)* mutant animals.

**Figure 3.**
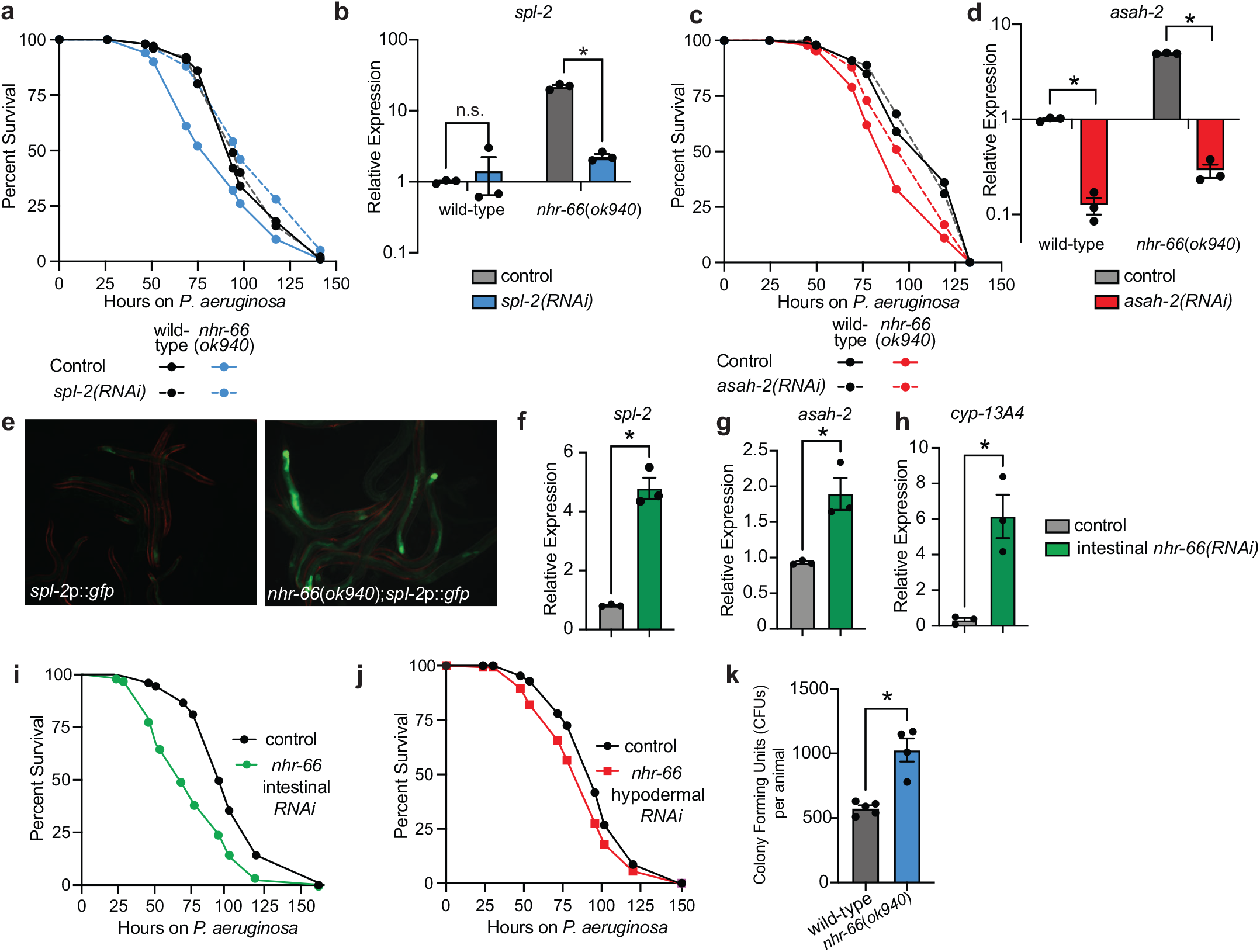
NHR-66 suppresses intestinal sphingolipid catabolism genes to promote resistance to *P. aeruginosa* infection. **A–D.** *C. elegans*-*P. aeruginosa* pathogenesis assay with *P. aeruginosa* and *C. elegans* of indicated genotypes at the L4 larval stage are shown (A and C). Differences in survival between *nhr-66(ok940*) mutants and the other genotypes in A and C are significant (*p*<0.05, log-rank test). Data representative of three biological replicates (*n*=3). qRT-PCR analysis of *spl-2* (B) and *asah-2* (D) genes in indicated genetic backgrounds. Data are the average of biological replicates with error bars showing SEM (n=3). * equals p<0.05 (two-way ANOVA with Tukey’s multiple comparisons test). **E.** Images of *C. elegans spl-2*p*::gfp* animals in wild-type and *nhr-66(ok940)* animals are shown. **F–H.** qRT-PCR analysis of the indicated genes in the indicated genetic backgrounds, as described in B and D. (n=3) **I and J.** *C. elegans – P. aeruginosa* pathogenesis assays as described above in A and C, except with the indicated genotypes. The difference between the two genotypes in I and J are significant (*p*<0.05, log-rank test). Data are representative of two biological replicates (n=2). **K.** *P. aeruginosa*, isolated from the intestines of animals with the indicated genotypes, were quantified after 24 hours of bacterial infection. Data are colony-forming units (CFU) of *P. aeruginosa* and are presented as the average of 4 separate biological replicates, with each replicate containing 10-11 animals. *equals p<0.05 (Student’s unpaired t-test). Sample sizes, mean lifespan and p-values for each replicate of the pathogenesis assays in this figure are shown in Table S5. Other source data for this figure is in Table S6. Please see also Figure S3.

We next asked in which tissue NHR-66 functions to repress sphingolipid catabolism and promote pathogen resistance. We generated a transcriptional reporter for the NHR-66 target, *spl-2*, by fusing its promoter to *gfp* (*spl-2*p::*gfp*). In the *nhr-66(ok940)* mutant, *spl-2*p::*gfp* was strongly induced and expressed in the intestinal epithelium (Fig. 3E). Consistent with these data, RNAi-mediated knockdown of *nhr-66* specifically in the intestine, using a *C. elegans* strain engineered to perform RNAi selectively in this tissue, increased the expression of *spl-2* (Fig. 3F), as well as two other NHR-66 targets [*asah-2* (Fig. 3G) and *cyp-13A4* (Fig. 3H)]. In addition, knockdown of *nhr-66* in the intestine rendered *C. elegans* hypersusceptible to *P. aeruginosa* infection (Fig. 3I). However, knockdown of *nhr-66* using a strain engineered to perform RNAi only in the hypodermis conferred only a subtle hypersusceptibility to pathogen infection (Fig. 3J). Of note, we previously tested the efficiency of RNAi in these engineered strains and found that *C. elegans* MGH167, the strain used for intestinal-specific RNAi, performs RNAi in the intestine, and not in the hypodermis, body wall muscle or germline^50^. Conversely, RNAi in *C. elegans* JM43, the strain used for hypodermal-specific RNAi, is still able to perform RNAi in the intestine, body wall muscle and germline, but not as efficiently as in the hypodermis^50^. Thus, in the *C. elegans* JM43 background, “off-target” RNAi-mediated knockdown of *nhr-66* in the intestine may account for the observed subtle enhancement of susceptibility to bacterial infection (Fig. 3J). Additionally, we observed that *C. elegans nhr-66(ok940)* mutants accumulated more *P. aeruginosa* in their intestines than wild-type animals, consistent with the enhanced susceptibility of this mutant to pathogen infection (Fig. 3K). Together, these data indicate that NHR-66 functions in the intestine to suppress the transcription of sphingolipid catabolism genes and promote pathogen resistance.

### Sphingolipid degradation in *nhr-66(ok940*) mutants compromises survival during pathogen infection

The NHR-66 targets, *spl-2* and *asah-2* are involved in multiple steps of the sphingolipid degradation pathway (Fig. 4A). To determine if NHR-66 regulates sphingolipid breakdown in *C. elegans*, we quantified sphingolipid levels in wild-type and *nhr-66(ok940)* mutants using high-performance liquid chromatography with tandem mass spectrometry (HPLC-MS/MS) (Fig. 4B-4E). We quantified 17 carbon and dihydro-17 carbon sphingolipids, which are the species produced in nematodes^51–53^. In metazoans, sphingolipids are broken down in cells by enzymes that are strongly conserved throughout evolution^54^. Sphingosine phosphate lyase (*C. elegans spl-1* and *spl-2*) and N-acylsphingosine amidohydrolase (also known as acid ceramidase, *C. elegans asah-1* and *asah-2*) function at sequential steps in the sphingosine catabolic pathway (Fig. 4A). SPL-1 and SPL-2 catalyze the final, irreversible step in the breakdown of sphingolipids to long-chain fatty acid aldehyde and phospho-ethanolamine (Fig. 4A). Thus, consistent with the high levels of SPL-2 expression in *C. elegans nhr-66*(*ok940*) animals, we observed that these mutants had significantly decreased levels of both sphingosine-1-phosphate and sphinganine-1-phosphate (Figs. 4A-C). The unphosphorylated species sphingosine and sphinganine were unchanged (Figs. 4B and 4C), although it is important to note that the levels of these lipids are maintained by both anabolic and catabolic enzymatic reactions (Figs. 4A). We also found that the levels of some, but not all, of the sphingolipid species dihydroceramides (Fig. 4D) and ceramides (Fig. 4E) were lower in *nhr-66(ok940)* mutants, consistent with their enhanced catabolism in this genetic background. In addition, NHR-66 does not regulate the transcription, nor bind to the promoters of paralogs *spl-1* (Fig. S3A) or *asah-1* (Fig. S3B) Thus, NHR-66 regulates sphingolipid catabolism by repressing the transcription of *spl-2* and *asah-2*, specifically.

**Figure 4.**
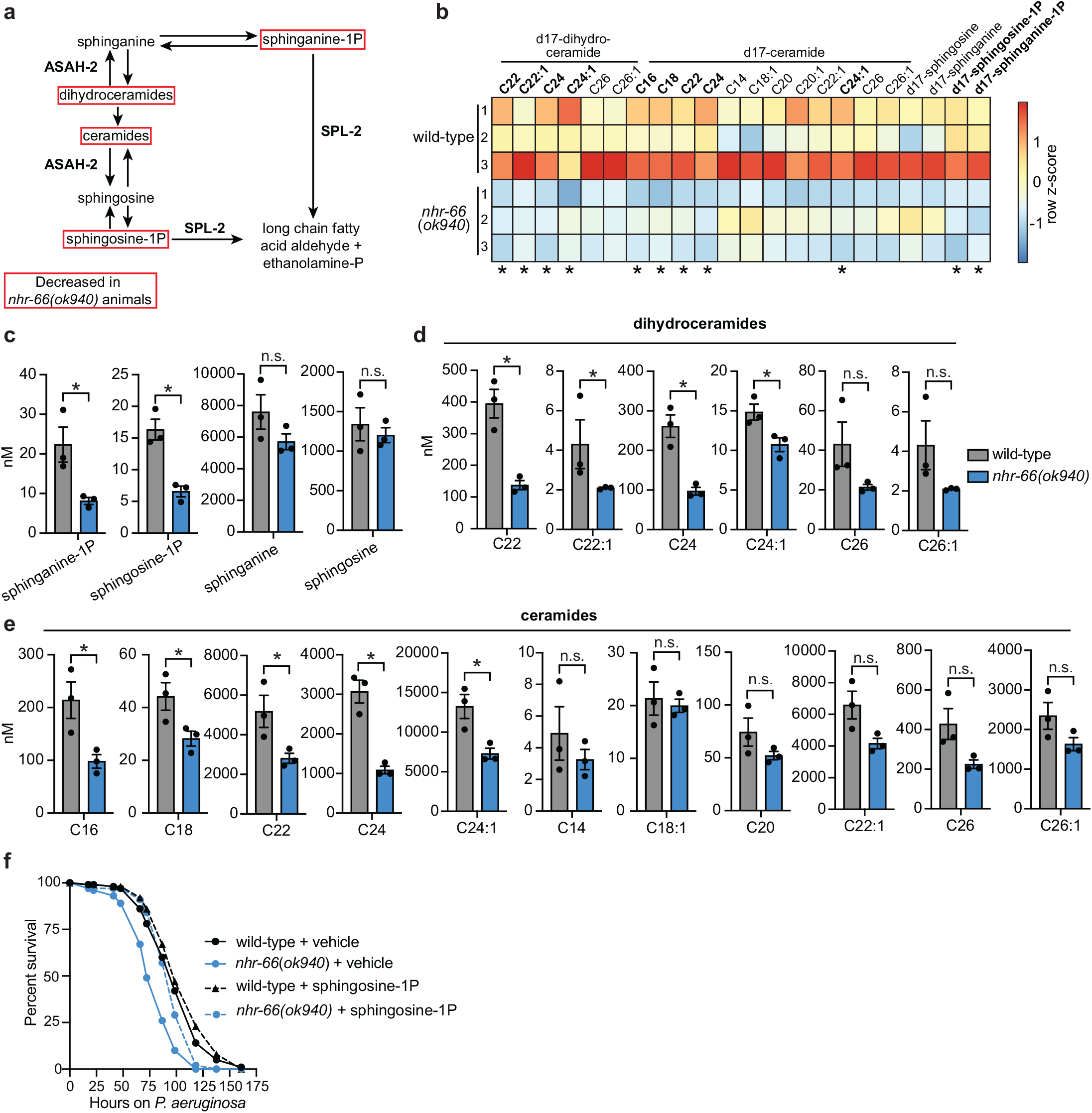
Sphingolipid degradation in *nhr-66(ok940*) mutants compromises survival during pathogen infection. **A.** Schematic of sphingolipid metabolism in *C. elegans*. **B.** Heatmap of HPLC-MS/MS data depicting the concentrations of d-17 ceramides and di-17 dihydroceramides normalized to total protein levels. The expression level was scaled in each condition by calculating a row z-score for each lipid species *p<0.05 (Student’s unpaired t-test) **C–E.** The concentrations of the indicated sphingolipids in wild-type versus *nhr-66*(*ok940*) animals as measured by HPLC-MS/MS. n=3 biological replicates, * p<0.05 (Student’s unpaired t-test), n.s. (not significant). **F.** *C. elegans-P. aeruginosa* pathogenesis assay with *C. elegans* of indicated genotypes at the L4 larval stage are shown. The difference in survival between *nhr-66(ok940*) mutants and other conditions is significant (p<0.05, log-rank test). Data representative of three biological replicates (n=3). Sample sizes, mean lifespan, and *p*-values for each replicate of this pathogenesis assay are in Table S5. The source data for this figure is in Table S6.

Intriguingly, supplementation of sphingosine-1-phosphate, a sphingolipid that is excessively catabolized in the *nhr-66(ok940)* background (Fig. 4C), partially rescued the hypersusceptibility of *nhr-66(ok940)* mutants during *P. aeruginosa* infection (Fig. 4F). These data indicate that hypersusceptibility of *nhr-66(ok940)* mutants to *P. aeruginosa* infection is secondary to excessive degradation of sphingolipids in this mutant background. It is important to note that sphingosine-1-phosphate can be rapidly converted to other sphingolipids in the cell (Fig. 4A). Thus, it is not possible to attribute the enhanced susceptibility of *nhr-66(ok940)* solely to reduced levels of sphingosine-1-phosphate.

In summary, transcriptional de-repression of the sphingolipid catabolism genes *spl-2* and *asah-2* in the *nhr-66(ok940)* mutant background accelerated the breakdown of sphingolipids in a manner that compromised the ability of *C. elegans* to survive bacterial infection. Thus, transcriptional control of sphingolipid catabolism by NHR-66 is necessary for pathogen resistance in *C. elegans*.

### NHR-66 and NHR-49 cooperate to promote pathogen resistance

NHR-66 physically interacts with another NHR, the *C. elegans* Peroxisome Proliferator-Activated Receptor (PPAR) homolog (NHR-49), a partnership that has been implicated in the regulation of sphingolipid catabolism genes^49^. NHR-49 is a master regulator of lipid metabolism in *C. elegans* that is also required for survival during pathogen infection^37,38,49,55^. We therefore hypothesized that NHR-49 functions with NHR-66 to promote resistance to *P. aeruginosa*. RNAi-mediated knockdown of *nhr-49* in the *nhr-66(ok940)* mutant background enhanced the susceptibility to pathogen-mediated killing to the same degree as *nhr-49(RNAi)* animals, suggesting that these NHRs cooperate to promote pathogen resistance (Fig. 5A). Consistent with these data, and a previous study^56^, we found that *nhr-49* and *nhr-66* regulated a significant number of genes in common (Fig. 5B), a group that is strongly enriched for metabolism and stress response genes (Fig. 5C). Notably, 13 of the 83 genes that are regulated in common by *nhr-49* and *nhr-66* were also among the 18 genes whose promoters were directly bound by NHR-66 in the ChIP-seq experiment (Fig. 2A). We examined the transcription of three of these genes (*spl-2, asah-2*, and *cyp-13A4*) and observed an additive effect of *nhr-49(RNAi)* and *nhr-66(ok940)* on the expression of *spl-2* (Fig. 5D) and *cyp-13A4* (Fig. S4), but not *asah-2* (Fig. 5E). Finally, we asked if the hypersusceptibility of *nhr-49* loss-of-function mutants was attributable to the hyper-induction of sphingolipid catabolism genes in this genetic background, as we observed in our studies of *nhr-66* (Fig. 3A-3D). Notably, RNAi of *spl-2* (Fig. 5F) or *asah-2* (Fig. 5G) did not rescue the enhanced susceptibly of *nhr-49(nr2041)* mutants to *P. aeruginosa* infection. These data are consistent with previous work, which has established that NHR-49 regulates multiple aspects of host metabolism that are individually important for survival during pathogen infection^37–39,49,55^. In summary, we conclude that NHR-66 and NHR-49 cooperate to regulate sphingolipid catabolism genes and promote pathogen resistance.

**Figure 5.**
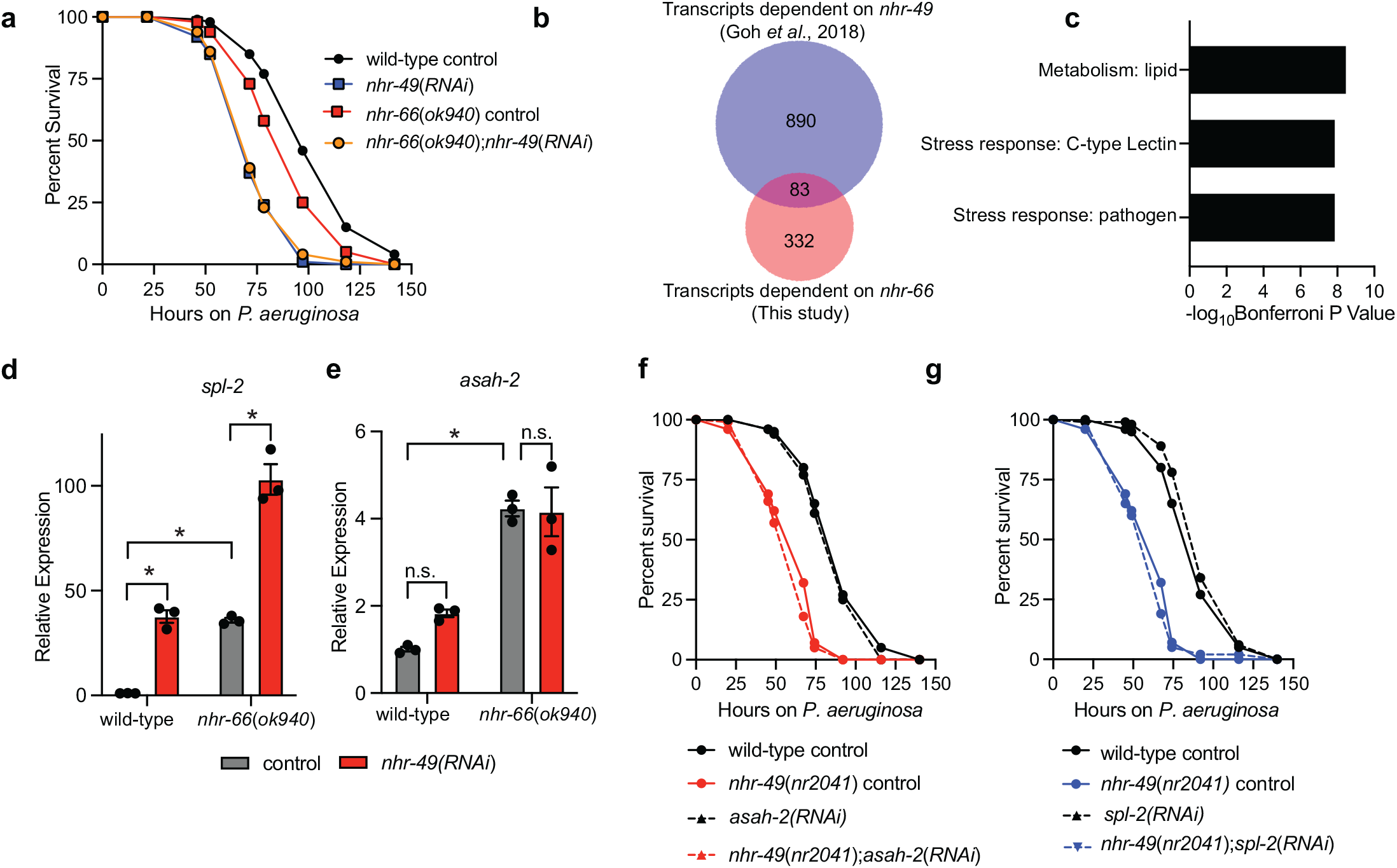
NHR-66 and NHR-49 cooperate to promote pathogen resistance. **A**. *C. elegans-P. aeruginosa* pathogenesis assay with *C. elegans* of indicated genotypes at the L4 larval stage are shown. Data are representative of three trials. The difference in survival between *nhr-66(ok940*) mutants and *nhr-66(ok940*); *nhr-49(RNAi)* animals is not significant. The other comparisons are significant (*p*<0.05, log-rank test). Data are representative of three biological replicates (n=3). **B.** Venn diagram showing the overlap between genes that are differentially regulated by *nhr-66* (this study) and those that are differentially regulated by *nhr-49*^56^. The overlap between these datasets is significant ( hypergeometric *p*-value = 7.9 x 10^-26^). See Table S4. **C.** Gene enrichment analysis using wormcat^80^ for the genes whose transcription are regulated by both *nhr-49* and *nhr-66* is shown. The three most significantly enriched categories are shown, reported as the log_10_ transformation of the *p*-value for the enrichment of each category. **D and E.** qRT-PCR analysis of the indicated genes in wild-type and *nhr-66(ok940)* mutant animals. Data are the average of three independent replicates with error bars representing SEM. Data are presented as the value relative to the average expression from all replicates of the indicated gene in the baseline condition (wild-type animals exposed to control). * *p*<0.05 (two-way ANOVA with Tukey’s multiple comparisons test). (F and G). *C. elegans-P. aeruginosa* pathogenesis assay with *C. elegans* of indicated genotypes at the L4 larval stage. Difference between *asah-2(RNAi)* (F) or *spl-2(RNAi)* (G) and control RNAi is not significant in wild-type animals but is significant in two out of three trials in *nhr-49(nr2041)* animals. Difference in survival between *nhr-49(nr2041)* and wild-type animals is significant (p<0.05, log rank). Data are representative of three biological replicates (n=3). Sample sizes, mean lifespan, and *p*-values for each replicate of the pathogenesis assays in this figure are shown in Table S5. Other source data for this figure is in Table S6.

### Regulation of sphingolipid catabolism by NHR-66 supports basal, rather than pathogen-induced, host defenses

*C. elegans*, like other metazoans, coordinate pathogen-inducible immune defenses through mechanisms that detect the pathogen itself or the effects of their secreted toxins. In addition, innate or basal defenses are also required for nematodes to survive challenges from infectious pathogens. Examples in this latter category include structural features of the cell (*e.g.*, membranes), the function of organelles, and micronutrients, such as cholesterol. To determine whether the regulation of sphingolipid catabolism by NHR-66 supports pathogen-induced or basal host defenses, we used HPLC-MS/MS to quantify sphingolipid levels in wild-type animals during infection with *P. aeruginosa*. None of the eleven NHR-66-dependent sphingolipids (Fig. 4) were upregulated in wild-type animals during infection with *P. aeruginosa*, as might be expected if these sphingolipids were coordinating an inducible response to pathogen infection (Fig. 6A). In addition, the transcription of *nhr-66* itself did not change in wild-type animals during *P. aeruginosa* infection (Fig. 6B).

**Figure 6.**
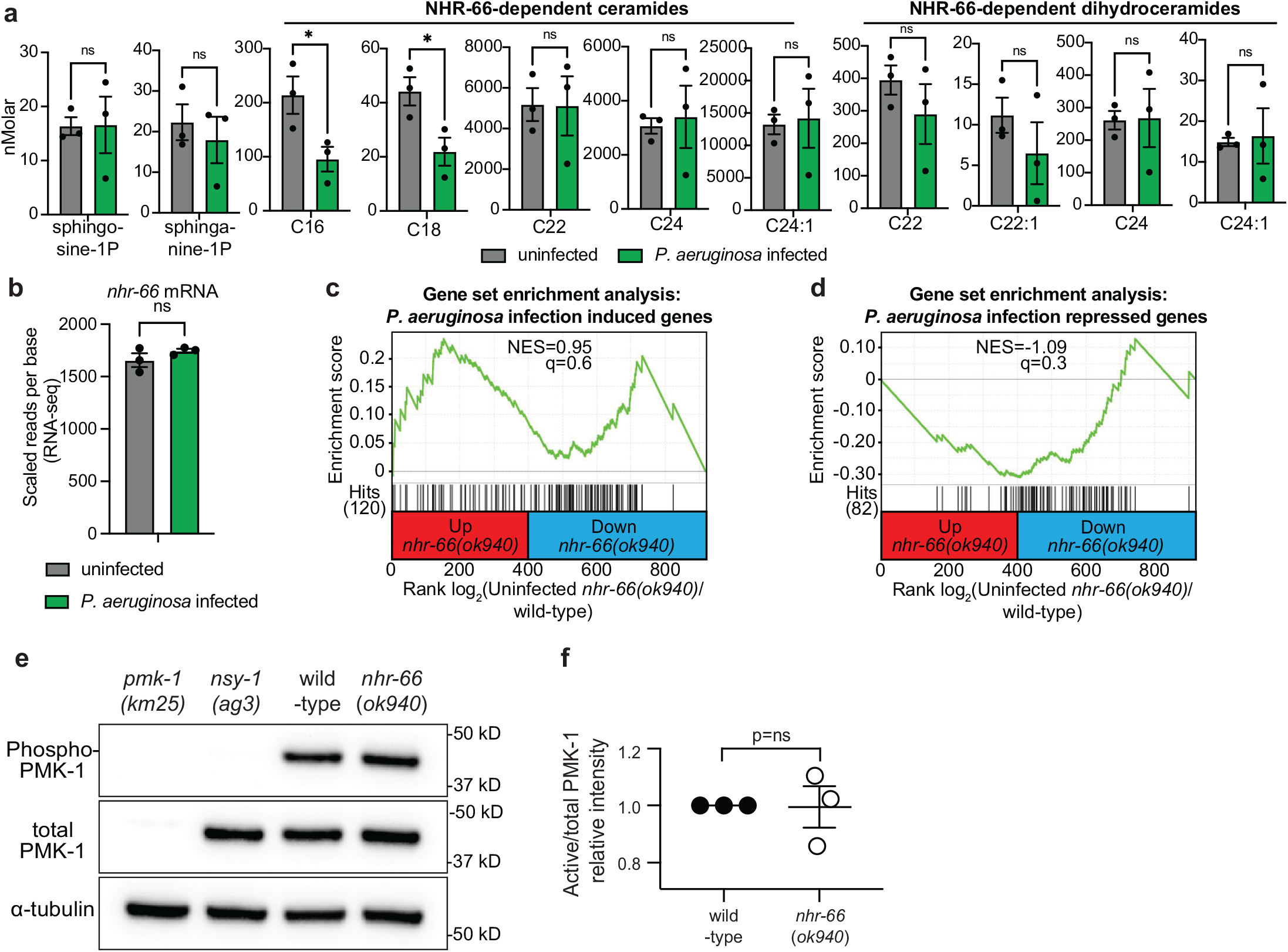
Regulation of sphingolipid catabolism by NHR-66 supports basal, rather than pathogen-induced, host defenses. **A**. The concentrations of the indicated sphingolipids in wild-type animals either during infection with *P. aeruginosa* or in uninfected controls, as measured by HPLC-MS/MS. Data are the average of three biological replicates with error bars equal to SEM, * *p*<0.05 (Student’s unpaired t-test) (n=3). **B.** RNA-seq data of the *nhr-66* transcript in wild-type animals either during infection with *P. aeruginosa* or in uninfected controls. Data are the average of three biological replicates with error bars equal to SEM. RNAseq data of *C. elegans* infected with *P. aeruginosa* was previously published.^41^ **C and D.** Gene set enrichment analysis (GSEA) for genes induced **(C)** or repressed **(D)** in *P. aeruginosa* infected animals in the RNA-seq of *nhr-66* dependent genes. In **(C)** and **(D)**, fold change in the expression of the significantly differentially expressed genes (q<0.01) in *nhr-66(ok940)* mutant animals compared to wild-type animals are ranked from higher expression (red) to lower expression (blue). Normalized enrichment score (NES) and q-value are indicated. Genes induced by either condition and found in the *nhr-66(ok940)* transcriptional profile are indicated by hit number in the left margin and black lines. **E.** Immunoblot analysis of lysates from the indicated genotypes probed with antibodies targeting the doubly phosphorylated TGY epitope in phosphorylated PMK-1 (phospho-PMK-1), total PMK-1 protein (total PMK-1), and tubulin (α-tubulin). *nsy-1(ag3)* and *pmk-1(km25)* loss-of-function mutants are the controls, which confirm the specificity of the phospho-PMK-1 probing. **F.** The band intensities of three biological replicates of the Western blot shown in (E) were quantified. Error bars reflect SEM. *equals *p*<0.05 (Student’s unpaired t-test). n.s. (not significant). Other source data for this figure is in Table S6.

We next asked if NHR-66 indirectly regulates the transcription of innate immune effectors to promote pathogen resistance (Figs. 6C and 6D). However, pathogen-response genes were not enriched among the genes regulated by *nhr-66*. Specifically, genes that are either upregulated (Fig. 6C) or downregulated (Fig. 6D) in wild-type animals during *P. aeruginosa* infection were not overrepresented among *nhr-66*-dependent genes. We also observed that *nhr-66* does not regulate the p38 mitogen-activated protein (MAP) kinase PMK-1 innate immune pathway. The levels of active, phosphorylated p38 PMK-1 were unchanged in *nhr-66*(*ok940*) mutants compared to wild-type animals (Figs. 6E and 6F).

Importantly, supplementation of sphingosine-1-phosphate did not appreciably extend the lifespan of wild-type animals during pathogen infection, with a subtle lifespan extension observed in only 2 of 3 biological replicates (Fig. 4F), arguing that sphingosine-1-phosphate or a derivative does not activate inducible immune defenses.

We therefore conclude that transcriptional control of sphingolipid catabolism by NHR-66 is required for basal, but not pathogen-inducible, host defense.

## DISCUSSION

Sphingolipids play critical roles in cellular physiology. Here we demonstrate that sphingolipid breakdown is regulated at the transcriptional level. We show that *C. elegans* NHR-66 directly regulates sphingolipid catabolism by repressing the transcription of two enzymes, sphingosine phosphate lyase (SPL-2) and acid ceramidase (ASAH-2). Transcriptional repression of these catabolic enzymes by NHR-66 controls the levels of sphingosine-1-phosphate, sphinganine-1-phosphate, and specific ceramides. Control of sphingolipid catabolism, in general, and *spl-2* and *asah-2* regulation specifically, is physiologically important and required for host survival during pathogen challenge. These data define an immunometabolic axis that is necessary for pathogen resistance (Fig. 7).

**Figure 7.**
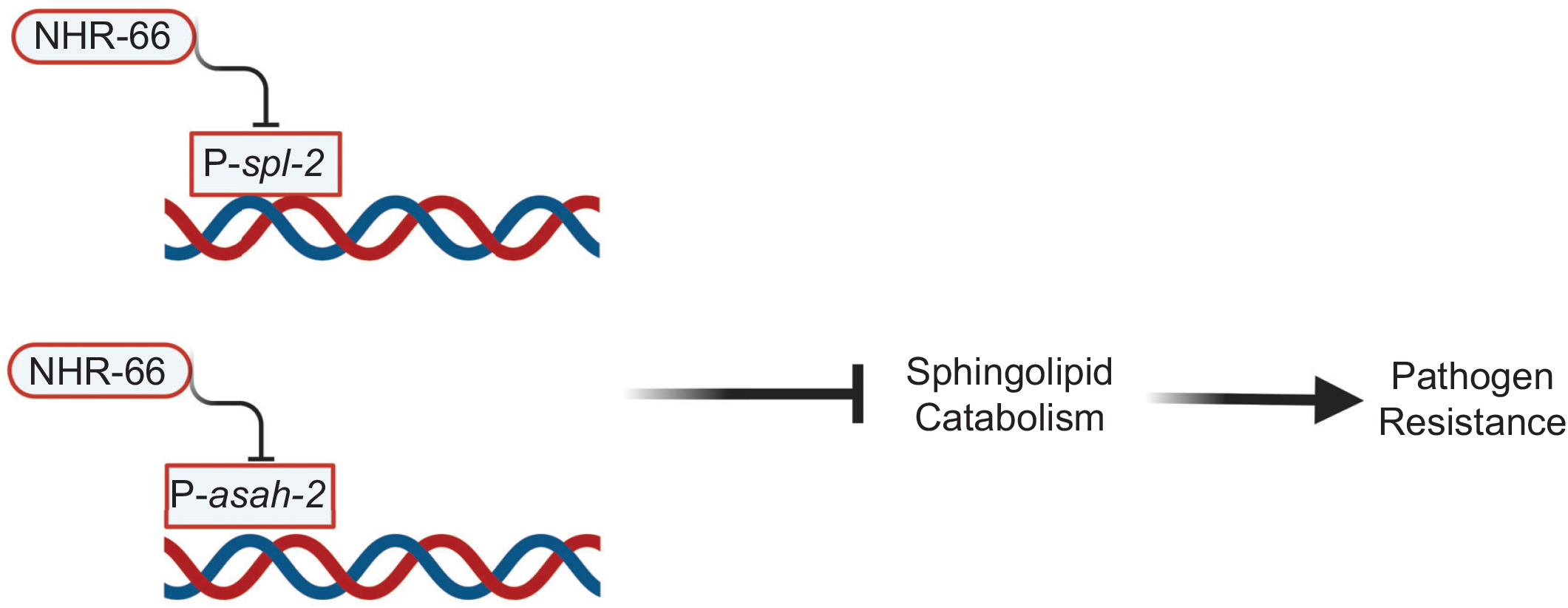
Transcriptional suppression of sphingolipid catabolism controls pathogen resistance in *C. elegans*.

Studies in *C. elegans* have revealed metabolic requirements for host survival during pathogen infection^45^. For example, the monounsaturated fatty acid oleate and the methyl donor S-adenosylmethionine promote innate immunity and pathogen resistance^39,40^. Likewise, insulin/insulin-like growth factor (IGF) signaling, through the insulin/IGF-1 transmembrane receptor homolog DAF-2, integrates nutritional cues and fat utilization to enable survival during bacterial infection^57^. In addition, mobilization of somatic fat to the germline is a protective response that ensures reproductive fidelity following pathogen challenge^58^. We also previously found that *C. elegans*, which encounter environments scarce in cholesterol, are more susceptible to pathogen-mediated killing and respond by priming innate immune signaling to anticipate threats from virulent bacteria^41^. Thus, a major emerging theme is the utility of the *C. elegans* pathogenesis assays to identify and characterize specific mechanisms of immunometabolism, studies that can be complicated or more challenging in other genetic systems.

Consistent with the important role of sphingolipids in host defense, a previous study found that mutation of the sphingosine kinase *sphk-1* and two putative sphingosine-1-phosphate transporters compromised host survival during *P. aeruginosa* infection^59^. However, it is not known how excess sphingolipid catabolism in the *nhr-66* mutants compromises host survival during pathogen challenge. Our data suggest that sphingolipids are required for basal mechanisms of host defense, but not pathogen-inducible responses, such as those coordinated by soluble signals. This distinction is important because sphingolipids as a group support diverse cellular processes with described functions both as bioactive signaling molecules and key components of cell and organelle membranes. We hypothesize that proper sphingolipid catabolism is important for maintaining membrane integrity, which is required for resistance against pathogens^12,60,61^. Sphingolipids are also important for mitochondrial function, which may be compromised in the *nhr-66* loss-of-function mutants^49,62^. Finally, the pathogen itself may exploit changes in host sphingolipid composition to promote its own virulence^63,64^.

*C. elegans* consume bacteria as their source of nutrition and live in habitats where both infectious pathogens and non-infectious microbes are abundant. Thus, identifying infectious pathogens is particularly challenging for nematodes. Intriguingly, *C. elegans* lost canonical mechanisms of pattern recognition [*e.g.,* Toll-like receptor (TLR) signaling] in evolution, perhaps because TLR ligands (pathogen-associated molecular patterns) are ubiquitous in their environment and are thus insufficient to identify disease-causing pathogens specifically. In this context, we previously proposed that the dramatic expansion of the NHR family in *C. elegans* was fueled, at least in part, by their roles in pathogen detection, immune activation, and immunometabolism^36,65^. In support of this hypothesis, we found that *C. elegans* NHR-86 senses a pathogen-derived metabolite to assess the relative threat of virulent bacteria in the environment and activate innate immunity — a non-canonical mechanism of bacterial pattern recognition^36^. As ligand-gated transcription factors, NHRs are also uniquely positioned to adapt host metabolism to survive challenges from infectious pathogens. In *C. elegans*, for example, NHR-49, the PPAR⍺ homolog, is a master regulator of lipid homeostasis that regulates a wide variety of genes involved in the synthesis and utilization of fatty acids and is required to survive pathogen challenge^37,38,48,49^. We observed here that NHR-49 cooperates with NHR-66 to regulate sphingolipid catabolism genes and survival during *P. aeruginosa* infection. These data support a previous study, which found that NHR-49 establishes functional partnerships with NHR-66 and other NHRs to regulate specific metabolic outputs, including genes involved in sphingolipid metabolism^49^. In this context, the ligands of NHR-66 or NHR-49 are not known. It is interesting to note that NHR-66 regulates its own transcription, suggesting that it functions in a feedback loop. Of note, we also report the identification of *nhr-32* from a comprehensive RNAi screen, which is required to resist *P. aeruginosa* infection. We hypothesize that examining the role of diverse pathogens in *C. elegans* will identify a different set of NHRs that survey for virulence-associated metabolites from the pathogen and program adaptive and pathogen-specific changes in host metabolism.

## ACKNOWLEDGEMENTS

The authors thank Melanie Trombly for critical reading of the manuscript. The authors also thank Gordon Worley, Eduardo Torres, Ryan Palumbo, and Jason Pierce for valuable discussions. This research was supported by R01 AI130289 (to R.P.W.), R01 AI159159 (to R.P.W.), a diversity supplement to R01 AI159159 (to M.A.N. and R.P.W.), R21 AI163430 (to R.P.W.), F30 AI150127 (to N.D.P.), F30 DK127690 (to S.Y.T.), T32 AI132152 (to N.D.P.), T32 AI095213 (to S.Y.T.), and T32 GM107000 (to M.A.N., N.D.P. and S.Y.T.). Some strains were provided by the *Caenorhabditis* Genetics Center, which is funded by the NIH Office of Research Infrastructure Programs (P40 OD010440).

## AUTHOR CONTRIBUTIONS

Conceptualization: MAN, NDP, RPW; Methodology: MAN, NDP, RPW; Investigation: MAN, NDP, JES, PL, ALP, SYT, KAW, CET; Visualization: MAN, NDP; Funding acquisition: RPW; Project administration: RPW; Supervision: RPW; Writing – original draft, MAN, RPW; Writing – review & editing, MAN, NDP, RPW.

## COMPETETING INTERESTS

The authors declare no competing interests.

## MATERIALS AND METHODS

### *C. elegans* and bacterial strains

The previously published *C. elegans* strains used in this study are: N2 Bristol (wild-type)^66^, *nhr-49*(*nr2041*)^55^, *nhr-8(hd117)*^30^*, nhr-8(ok186)*^67^, VC1727 *nhr-32(gk825)*^68^, VC1692 *nhr-32(gk810)*^68^, KU25 *pmk-1(km25)*^69^*, nsy-1(ag3)*^69^, MGH167 *sid-1(qt9); alxIs9 [VHA-6*p*::SID-1::SL2::GFP]*^70^, JM43 *(rde-1(ne219); Is[wrt-2*p*::rde-1]*^71^, and *nhr-66*(*ok940*)^49^.

The *C. elegans* strains that were developed for this study are: RPW 451 *nhr-66(ums70[3xFLAG::NHR-66]),* RPW 257 *nhr-66*(*ok940*);Fosmid line 1, RPW 257 *nhr-66*(*ok940*);Fosmid line 2, *spl-2*p::gfp, *nhr-66*(*ok940*);*spl-2*p::gfp, RPW 436 *nhr-32(gk810);umsEx89(nhr-32p::FL-nhr32::myo-3::mCherry)* line 1; RPW 437 *nhr-32(gk810);umsEx90nhr-32p::FL-nhr32::myo-3::mCherry)* line 2, and RPW 280 *nhr-8(hd117);umsEx42(nhr-8p::nhr-8, myo-2p::mCherry) Line 1*, RPW 281 *nhr-8(hd117);umsEx42(nhr-8p::nhr-8, myo-2p::mCherry) Line 2. P. aeruginosa* strain PA14 was used for all studies^72^.

### *C. elegans* bacterial infection and other assays

*P. aeruginosa* “slow-killing” pathogenesis experiments were performed as previously described^50,73^. The wild-type control for these experiments is N2. Briefly, *P. aeruginosa* PA14 was inoculated into 3 ml of Luria-Bertani (LB) medium and allowed to incubate at 37 °C for 16 hours. Subsequently, 10 μl of this culture was spread onto 35-mm tissue culture plates containing 4 ml of slow-kill agar (0.35% peptone, 0.3% sodium chloride, 1.7% agar, 5 μg/ml cholesterol, 25 mM potassium phosphate, 1 mM magnesium sulfate, 1 mM calcium chloride). Of note, we used peptone from Fisher Scientific for these studies. Plates were incubated for 24 hours at 37 °C, and 24 hours at 25 °C. Twenty minutes before the start of the assay, 0.1 mg/ml 5-fluorodeoxyuridine was added to the medium to prevent progeny from hatching. For all pathogenesis assays that studied *C. elegans* with extrachromosomal arrays, control genotypes, which did not express the array, were obtained from siblings isolated from the same plates as nematodes that contained the array. *C. elegans* lifespan assays were conducted with animals grown on nematode growth media agar at 20 °C in the presence of 40 µg/mL 5-fluoro-2’-deoxyuridine. All pathogenesis and lifespan assays were conducted with nematodes at the L4 larval stage. Unless noted, three independent trials of each pathogenesis assay were performed. Sample sizes, mean lifespan, and *p*-values for all trials are shown in Table S5.

Colony-forming units of *P. aeruginosa* PA14 were counted in the intestine of *C. elegans* as previously described^50,74^. In brief, *C. elegans* were exposed to *P. aeruginosa* for 24 hours. Animals were then picked to NGM plates lacking bacteria and incubated for 10 minutes to remove external *P. aeruginosa*. Animals were then transferred to a second NGM plate after which 9–11 animals per replicate were collected, washed with M9 buffer containing 25 mM levamisole and 0.01% Triton X-100, and ground with 1.0 mm silicon carbide beads. CFUs were counted from serial dilutions of the lysate.

### RNAi screen for *nhr* genes required for pathogen resistance in *C. elegans*

A previously described library containing RNAi clones corresponding to 271 of 274 *nhr* genes in the *C*. *elegans* genome was used for this study^36,75^. *C. elegans* exposed to each of these RNAi clones were screened for survival on *P. aeruginosa* PA14 following the “slow-kill” protocol described above. In the primary screen, the survival of ∼50 animals on a single killing assay plate was assessed. RNAi clones that caused 0.75 fraction mean survival or greater compared to wild-type were selected and then screened with ∼150 animals on three different plates (∼50 animals per plate). In the secondary screen, confirmed hits were assayed for lifespan defects by exposing worms to the RNAi clones and transferring them at the L4 stage to non-pathogenic *E. coli* OP50. In the tertiary screen, three independent trials, each with 150 animals, were performed on hits from the secondary screen.

### Generation of transgenic *C. elegans* strains

A fosmid containing 31 kb of *C. elegans* DNA (IV:8,209,892..8,240,932), which includes the promoter and coding region for *nhr-*66, as well as for *serp-1.2*, *gpx-6*, *ocr-2*, and C07G1.7, was microinjected into the gonad of *C. elegans nhr-66(ok940)* mutants, and two independent lines that expressed the fosmid in extrachromosomal arrays were recovered. The fosmid was obtained from Source Biosciences. To generate NHR-66::FLAG animals, CRISPR-Cas12 or Cpf-1 editing with single-stranded oligodeoxynucleotide (ssODN) homolog-directed repair was used to knock-in a 3xFLAG sequence at the C-terminus of the *nhr-66* coding sequence, just upstream of the stop codon, as previously described^36,76,77^. The C-terminus was selected for the 3xFLAG knock-in to label all *nhr-66* isoforms. To generate *nhr-32* rescue lines, PCR was used to amplify the entire *nhr-32* locus. The resulting PCR product (10 ng/μL), the *myo-3*p*::mCherry* co-injection marker (15 ng/μL), and pUC19 vector (135 ng/ml) were microinjected into *nhr-32*(*gk810*) animals to generate two independent lines. The list of oligos used in this study is in Table S7.

### Chromatin immunoprecipitation sequencing and bioinformatics

Chromatin immunoprecipitation was performed with a strain containing a 3xFLAG-tagged NHR-66 protein (NHR-66::3x-FLAG) generated in this study, as described above. L4 synchronized hermaphrodite *C*. *elegans* (wild-type and transgenic NHR-66::3x-FLAG animals) were collected and washed with 4 °C M9 and phosphate-buffered saline to remove bacteria. Cross-linking of protein and DNA was performed in 1% formaldehyde for 10 minutes at room temperature. Cross-linking was quenched with 100 mM glycine, animals were washed in M9, and resuspended in lysis buffer (50 mM Hepes–KOH pH 7.5, 300 mM NaCl, 1 mM EDTA, 1% (v/v) Triton X-100, 0.1% (w/v) sodium deoxycholate, 0.5% (v/v) N-Lauroylsarcosine, and protease inhibitors). Lysates were then sonicated using a Bioruptor UCD-200 for 15 cycles (30s on, 30s off). Anti-flag antibody was pre-incubated with protein G Dynabeads (Invitrogen), and equal amounts of protein from lysate were incubated with 5 μg anti-FLAG antibody (Roche) overnight. 10% of lysate was removed for input. Immune complexes were collected with protein G Dynabeads, washed, and eluted from beads. Cross-links were reversed at 65 °C overnight and DNA fragments were purified with ethanol precipitation. qPCR was performed on input and immunoprecipitated samples using primers designed around the transcription start site. ChIP data are presented as percent input normalized to a random intragenic region on chromosome IV. Primers used for ChIP-PCR studies are in Table S7.

Deep sequencing of ChIP DNA was performed by Novogene. The raw sequencing data were first clipped for adaptor sequences and then mapped to the *C*. *elegans* genome (ce11, UC Santa Cruz) by the Burrows-Wheeler Aligner algorithm (BWA MEM, BWA version 0.7.15). The output SAM files were processed and sorted with the Picard tools. The output mapping files (BAM files) were filtered with SAMtools to remove any read that had a mapping quality less than 10 (SAMtools view–b–q 10 input.bam > output.bam). Peaks were determined using MACS version 2.1 with the no-model parameter. The final set of peaks were called if the difference in intensity values of samples had a significance level of *p*-value < 0.025. ChIP-sequence data is available in Table S3.

### Gene expression analyses and bioinformatics

Approximately 2,500 synchronized *C. elegans* of the indicated genotypes were grown to the L4 stage and harvested by washing with M9. For expression analysis of *C. elegans* genes during *P. aeruginosa* infection, animals at the L4 stage were transferred by washing to plates containing *E. coli* OP50 or *P*. *aeruginosa* PA14 lawns, prepared as described above. Animals were exposed for four hours and subsequently harvested by washing with M9. RNA was isolated using TriReagent (Ambion), column purified (Qiagen), and analyzed by 150 bp paired-end mRNA-sequencing using the BGI platform (BGI Group) with > 20 million reads per sample. Raw fastq reads were evaluated by FastQC (version 0.11.5), and clean reads were aligned to the *C. elegans* reference genome (WBcel235) and quantified using Kallisto (version 0.45.0)^78^. Differentially expressed genes were identified using Sleuth (version 0.30.0)^79^. Pearson correlation statistical analysis was performed using Prism 9.0. Biological process enrichment was identified using Wormcat^80,81^. Differential gene expression was defined as q less than 0.01. Differentially expressed genes from the RNA-seq experiment are available in Table S2.

For the qRT-PCR studies, RNA was reverse transcribed to cDNA using the iScript cDNA Synthesis Kit (Bio-Rad Laboratories, Inc.), amplified and detected using SYBR Green (Bio-Rad Laboratories, Inc.) and a CFX384 machine (Bio-Rad Laboratories, Inc.). The sequences of primers that were designed for this study are presented in Table S7. All values were normalized against the geometric mean of the control genes *snb-1* and *act-3*. Fold change was calculated using the Pfaffl method^82^.

### Ceramide and sphingolipid profiling

High-performance liquid chromatography with tandem mass spectrometry (HPLC-MS/MS) for quantification of sphingolipid species was performed in the Lipidomics Shared Resource Facility at the Medical University of South Carolina on a Vanquish UHPLC system coupled to a Quantum Access Max triple quadrupole mass spectrometer equipped with an ESI probe operating in the multiple reaction monitoring positive ion mode (Thermo Scientific). Chromatographic separations were obtained under a gradient elution on a C8 column using a mobile phase with ammonium formate, formic acid in water and methanol, as previously described^83^. Prior to analysis, samples underwent an ethyl acetate/isopropanol liquid-liquid extraction. Quantitative analyses of sphingolipids were based on eight-point calibration curves generated for each target analyte. The synthetic standards along with a set of internal standards were spiked into an artificial matrix; they were then subjected to an identical extraction procedure as the biological samples. These extracted standards were then analyzed with the samples by HPLC-MS/MS. Peaks for the target analyses and internal standards were recorded and processed using the instrument’s software. Plotting the analyte/internal standard peak area ratios against analyte concentrations was used to generate the sphingolipid-specific calibration curves. Any sphingolipids for which no standards were available were quantitated using the calibration curve of its closest counterpart. Sample sphingolipid levels were normalized to total protein and volume.

### Western blots

Protein lysates from *C. elegans* grown to the young L4 larval stage on *E. coli* OP50 on NGM agar were prepared as previously described^36,41^. Briefly, animals were washed three times with M9 buffer, and lysed by sonication in RIPA buffer plus protease and phosphatase inhibitor cocktail. Following centrifugation, protein was quantified from the supernatant of each sample using Bradford Reagent (Bio-Rad Laboratories, Inc.). Laemmli buffer (Bio-Rad Laboratories, Inc.) was added to a concentration of 1X and the total protein from each sample was resolved on NuPage 4–12% gels (Life Technologies), transferred to nitrocellulose membranes (Life Technologies), blocked with 5% fat-free milk in TBST and probed with a 1:1000 dilution of an antibody that recognizes the doubly-phosphorylated TGY motif of PMK-1 (Cell Signaling Technology) or a monoclonal anti-tubulin antibody (Sigma-Aldrich, Clone B-5-1-2). Horseradish peroxidase (HRP)-conjugated anti-rabbit (Cell Signaling Technology) and anti-mouse IgG secondary antibodies (Abcam) were used at a dilution of 1:10,000 to detect the primary antibodies following the addition of ECL reagents (Thermo Fisher Scientific), which were visualized using a ChemiDoc MP Imaging System (Bio-Rad Laboratories, Inc.).

### Microscopy and imaging

*C. elegans* were mounted onto 2% agarose pads, paralyzed with 10 mM levamisole (Sigma), and photographed using a Zeiss AXIO Imager Z2 microscope with a Zeiss Axiocam 506 mono camera and Zen 2.3 (Zeiss) software at 10X magnification.

### Quantification and statistical analyses

Differences in survival of *C. elegans* in the *P. aeruginosa* pathogenesis assays were determined with the log-rank test after survival curves were estimated for each group with the Kaplan-Meier method. OASIS 2 was used for these statistical analyses^84^. *P*-values were calculated in Prism 9 (GraphPad Software) using one-way ANOVA, unless otherwise indicated in the figure legend. Sample sizes, mean lifespan, and p-values for all trials are shown in Table S5.

### Data availability

The mRNA-seq and ChIP-seq datasets were deposited in the NCBI Gene Expression Omnibus, a publicly available database, using the Accession numbers GSE240425 and GSE240426, respectively. All other data is available in this manuscript. Tables S5 and S6 contain all source data and statistical tests used in this manuscript.

## FIGURE LEGENDS

**Supplementary Figure 1.**
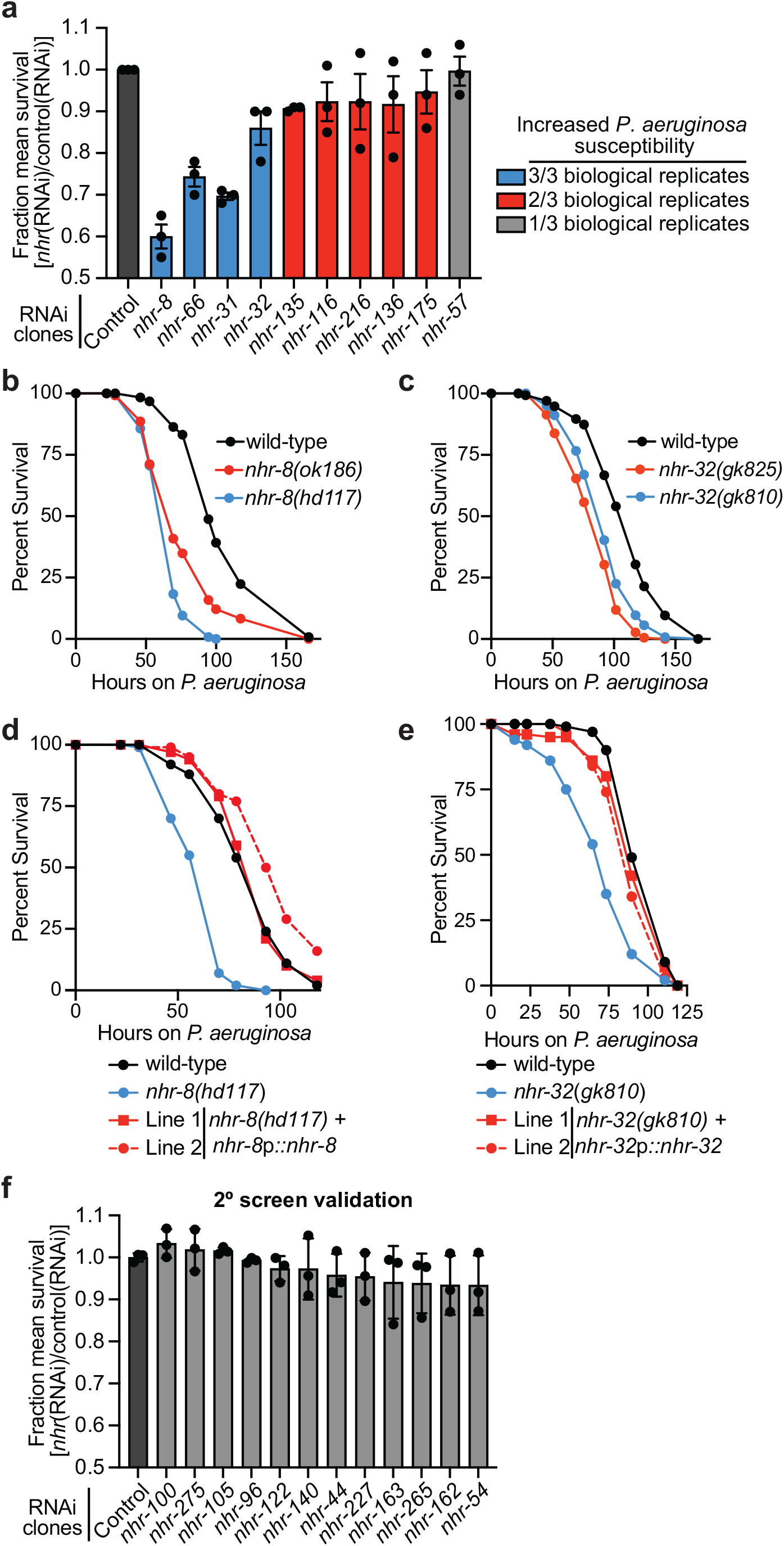
An RNAi screen identifies nuclear hormone receptors required for pathogen resistance in *C. elegans*. **A and F**. Fraction mean survival during *P. aeruginosa* infection of the indicated genotypes. See legend for Fig. 1C. **B-E.** *C. elegans-P. aeruginosa* pathogenesis assays, as described for Fig. 1E. Related to Fig. 1.

**Supplementary Figure 2.**
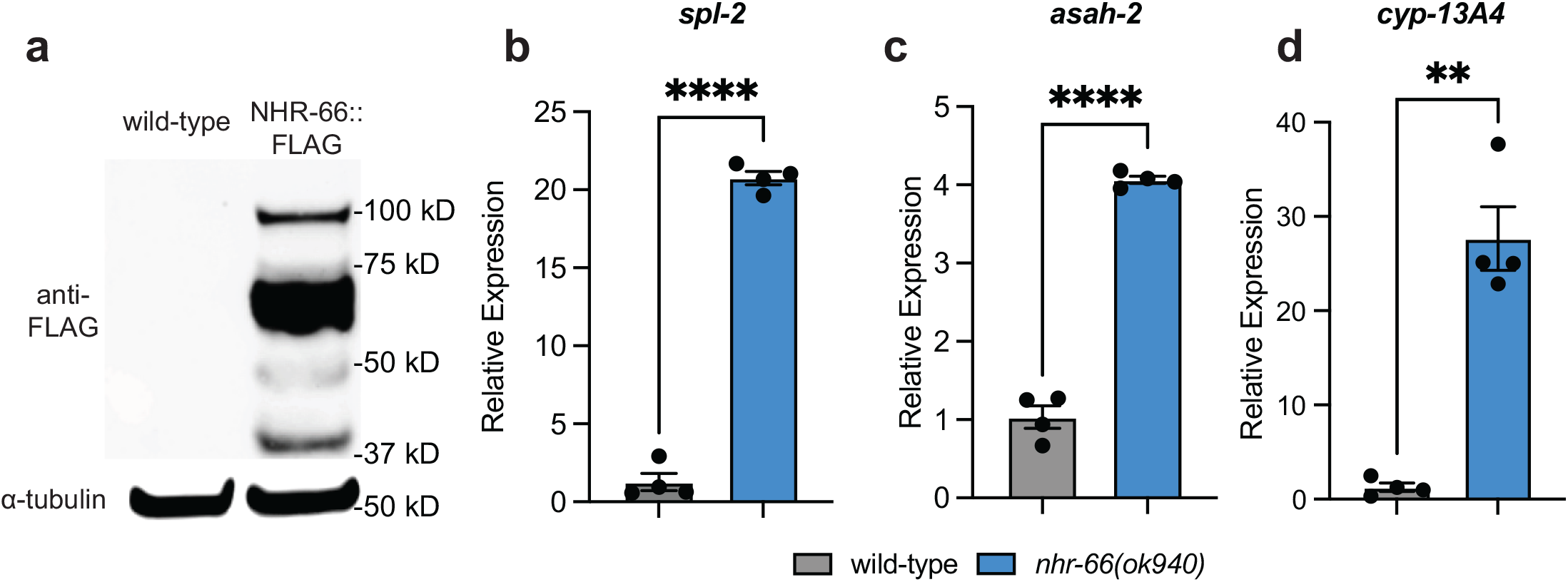
NHR-66 is a transcriptional repressor that binds to promoters. **A.** Western blot analysis of lysates from wild-type and NHR-66::3xFLAG animals probed with antibodies targeting the FLAG epitope (anti-FLAG) and α-tubulin (anti-tubulin). **B–D.** qRT-PCR analysis of the indicated genes in wild-type and *nhr-66(ok940)* mutant animals. Data are the average of four independent replicates with error bars representing SEM. * *p*<0.05 Student’s unpaired t-test). Source data for this figure is in Table S6. Related to Fig. 2.

**Supplementary Figure 3.**
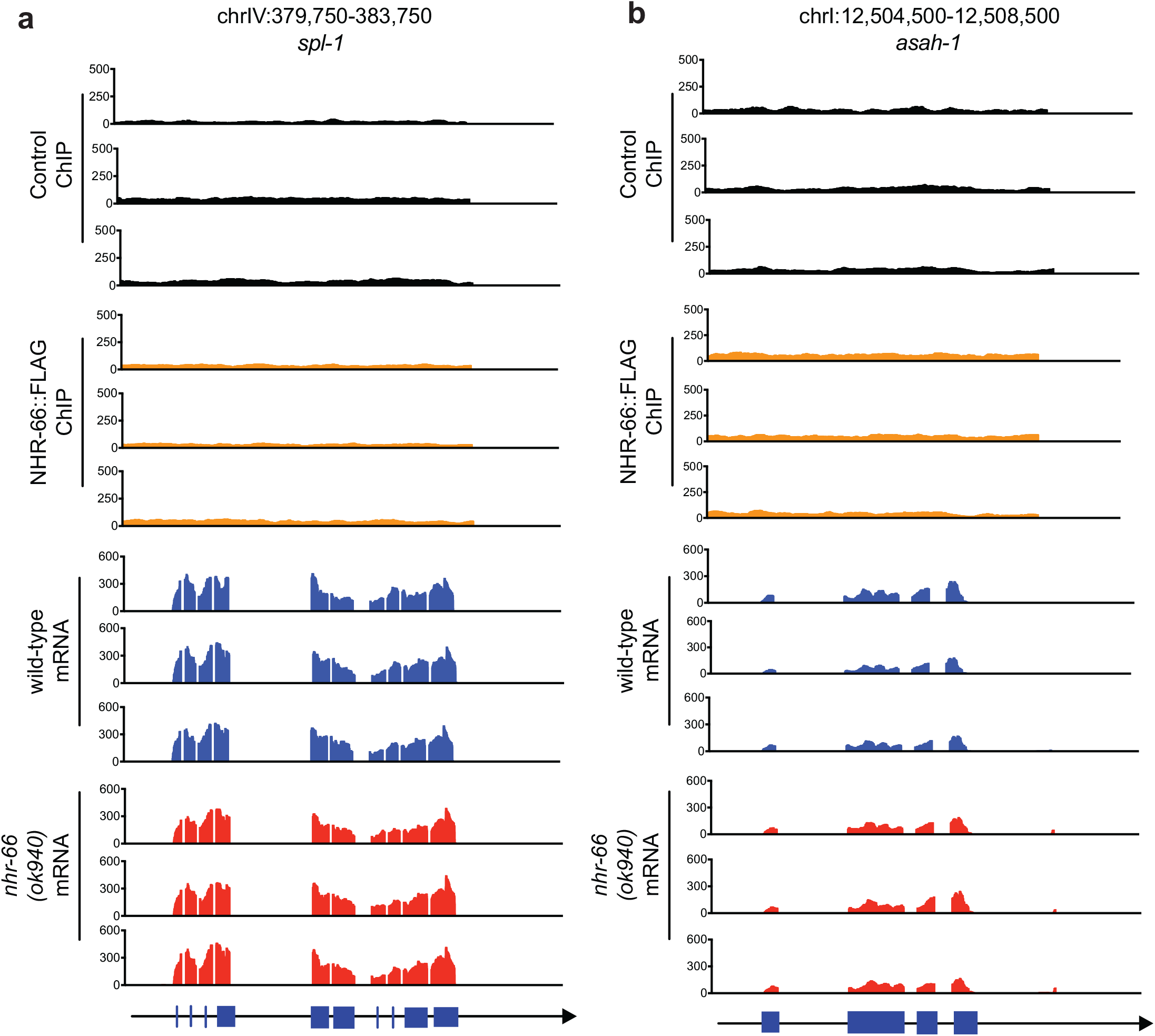
NHR-66-mediated repression of intestinal sphingolipid catabolism genes promotes pathogen resistance. **A-B.** ChIP-seq and mRNA-seq profiles from each of the three biological replicates are presented for *spl-1* (A), *and asah-1* (B). The y-axis is the number of reads (log_2_). A gene model shows the location of the exons (blue) of the indicated genes. Related to Fig. 3.

**Supplementary Figure 4.**
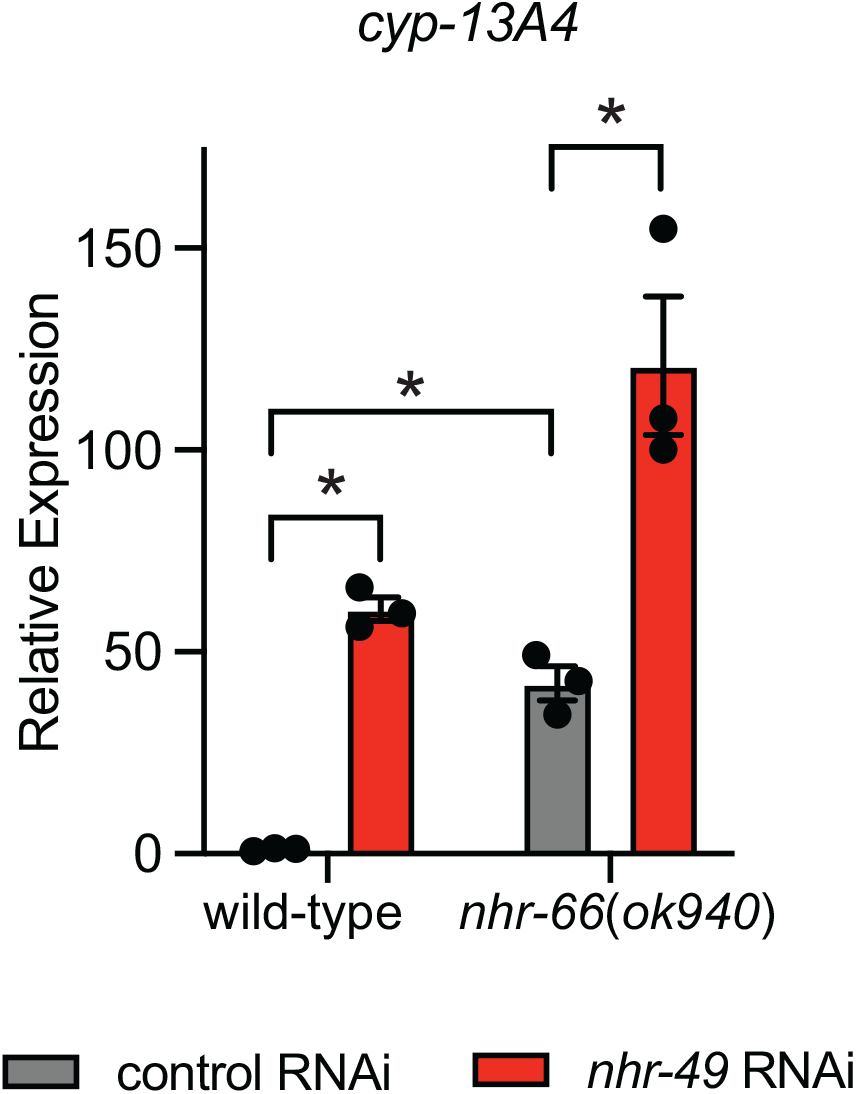
NHR-66 and NHR-49 cooperate to promote pathogen resistance. qRT-PCR data as described in Fig. 5D.

**Table S1. Source data for the RNAi screen for nuclear hormone receptor genes that are required for pathogen resistance in *C. elegans***. Related to Figure 1.

**Table S2. Data from RNA-seq experiment comparing wild-type to *nhr-66(ok940)* animals**. Related to Figure 2.

**Table S3. Data from ChIP-seq experiment to identify binding targets of NHR-66.** Related to Figure 2.

**Table S4. Overlap of genes regulated by both *nhr-66* and *nhr-49***. Related to Figure 5.

**Table S5. Sample sizes, mean lifespan, and p values for the *C. elegans* pathogenesis assays.** Related to Fig. 1E, Fig. 3A, Fig. 3C, Fig. 3I, Fig. 3J. Fig. 5A, Fig. 5G, Fig. 5H, Fig. S1C, Fig. S1D, and Fig. S1E.

**Table S6. Source data and statistical tests used for each figure and supplemental figure**. Related to Figs. 1-6, S1-S3.

**Table S7. Primer, crRNA guide sequences, and ssODN sequences used in this study**. Related to Materials and Methods.

